# Exploiting gene expression profiles of circulating extracellular vesicles for breast cancer detection

**DOI:** 10.1101/2024.05.09.593454

**Authors:** Aritra Gupta, Rosina Ahmed, Sanjit Agarwal, Geetashree Mukherjee, Kartiki V. Desai

## Abstract

**Background:** Liquid biopsy-based biomarkers offer several advantages since they are minimally invasive, can be useful in longitudinal monitoring of the disease and have higher patient compliance. We hypothesize that RNA content of circulating EVs differs in breast cancer patients and healthy women. EV RNAs may provide an opportunity to diagnose, detect subtypes and metastatic states.

**Experimental Design:** RNA-seq analysis and qRT-PCR from matched tumor biopsy, circulating EVs from breast cancer patients (EV-C) and healthy donors (EV-H) was performed to find genes that discriminate between these groups.

**Results:** EV-C to EV-H comparison yielded 320 DEGs (adjusted *p* value ≤0.05) enriched for cancer related pathways like Myc, Reactive oxygen species, and Angiogenesis. Allograft rejection and Interferon pathway genes were over-represented in the cancer group. Top 6 genes were validated by qRT-PCR in a validation cohort. 5 genes consistently and correctly classified cancer and healthy groups. An independent set of 374 and 640 DEGs could segregate ER positive/ER negative and non-metastatic versus metastatic samples, respectively. EVs from metastatic samples had higher variability in gene expression patterns whereas those from non-metastatic samples showed better correlation. Ability of 4 genes to classify metastases state was explored.

**Conclusion:** We report five EV RNAs that can be used to diagnose breast cancer in a subtype independent manner. Initial analysis indicates that EV RNA content differs based on subtype specificity and metastasis status.

## Introduction

Breast cancer is the most frequently occurring and a leading cause of cancer related deaths in women worldwide (https://www.wcrf.org/cancer-trends/breast-cancer-statistics). Generally, early detection of breast cancer leads to a favourable prognosis. Locally advanced cancers, local and distal metastases negatively impact survival, and account for more than 80% of cancer deaths (1). Molecular heterogeneity in breast cancer, development of drug resistance and clonal evolution during cancer treatment or progression also contribute to these outcomes. This implies that dynamic monitoring of tumor status in a longitudinal setting may facilitate improvement in treatment decisions. Repeated tissue biopsies during the duration of treatment are impractical, as they are invasive and may suffer from sampling biases. As a result, repeat samples for continuous assessment over the duration of treatment are rarely available (2). Recent advances show that diagnosis and real-time longitudinal monitoring of disease states may be achieved by non-invasive liquid biopsy approaches (2). Techniques such as circulating tumor cells (CTCs) and cell free DNA (cfDNA) can be employed, however their limiting concentrations in body fluids and the requirement for sophisticated infrastructure for isolation and purification, have imposed constraints on their use in longitudinal tumor surveillance (3–5).

Extracellular vesicles (EVs) including exosomes are 50-150 nm messengers carrying DNA/RNA, lipids and protein cargo enclosed by a lipid bilayer (6,7). They are released from most cells of the body and abundantly found in body fluids. Each day, most cells release 10^4^ particles in circulation, and they are a relatively stable source of their cargo (8). EVs deliver their contents both locally and distally via circulating blood, to elicit normal as well as pathophysiological responses in target cells. For example, tumor derived EVs transform non-cancerous cells, promote angiogenesis, and prepare pre-metastatic niche to increase metastatic spread (9). As EVs are capable of miRNA biogenesis, processing and delivery to affect target cell activities, several studies have concentrated on sampling EV miRNAs for tumor classification and discovery of novel biomarkers (10). Circulating EV miRNA and proteins are currently emerging as useful tools in diagnosis and prognosis of multiple cancers (11). Long non-coding RNAs and messenger RNAs (lncRNAs and mRNAs) carried by EVs have been well documented in several cell lines and their role in cancer is being studied (12). RNA profiles of EVs/exosomes are often phenocopies of their originating cells due to passive packing, but it is also reported that certain RNA transcripts are specifically enriched or excluded by as yet uncharacterized mechanisms (13). It can be envisaged that tumor derived EVs may differ from their normal cell counterparts in mRNA profiles, and transcriptomics of circulating EV RNA could be used to determine both the presence or absence of a tumor (diagnostic) and the response of the tumor to therapy. Further, identities of these mRNAs/lncRNAs delivered to local and distal sites will unravel molecular events participating in a pro-cancerous or pro-metastatic tumor microenvironment. However, robust protocols for transcriptome analysis of extremely low amounts of mRNAs carried by EVs are still being standardized, and only a handful of reports have characterized the transcriptome of patient-derived EVs in circulation (14). Here, RNA-seq analysis of circulating EV RNAs from breast cancer patient sera (EV-Cs), matched breast tumor biopsy samples and age-appropriate healthy control sera (EV-Hs) was undertaken. Our data indicates that patient EV RNA profiles substantially differ from healthy EVs suggesting that they may have diagnostic and disease monitoring potential. Further, our data indicates that additional application of EV RNA profiles may be useful in discriminating metastatic versus non-metastatic disease.

## Materials and Methods

### Sample collection

Core biopsies from breast cancer patients and matched blood samples (28 pairs) were collected at Tata Medical Center, Kolkata after Institutional review board approval an informed consent from patients (IRB EC/PHARMA/34/17). 3 paired samples of unknown clinical annotation were also included in this study. 14 samples blinded for their clinical status were used as validation cohort. Clinical details of the samples are mentioned in Supplementary table 1. Serum samples from seven healthy females (age 22-52 years) were used as healthy controls, 4 were subjected to RNA-sequencing.

### Purification and Analysis of EVs

Serum samples were centrifuged twice at 10,000 rpm for 10 minutes to remove any particulate matter. Exosome enriched EVs (hereafter EVs) were isolated from 500 µl serum using EXOQUICK (System Biosciences, CA) exosome precipitation method by standard protocols provided by the manufacturer. Nanoparticle tracking analysis (NTA) was used for quantification and sizing of EVs by Nanosight NS300 (Malvern Analytical Inc., West Borough, MA). For all runs, samples were injected and 60s video were captured thrice with camera level set at 7 and injection speed at 50 particles/S. Laser of 405 nm was set. Temperature was kept constant throughout the process. Size distribution and concentration of EVs were calculated by NTA software version 4.1. EVs from healthy controls are henceforth referred to as “EV-H” and EVs from cancer serum samples as “EV-C”. Western Blotting of EVs protein was performed using slot blot EXO-Array Exosome antibody check kit (System Biosciences, CA). Antibody slots for CD63, EpCAM, ANXA5, TSG101, GM130, FLOT1, ICAM, Alix and CD81 were present. Standard kit protocol was used for processing and development. For detection of chemiluminescence of the blot Chemidoc (Bio Rad, USA) was used.

### RNA sequencing and transcriptome analysis

Since EVs contain extremely low levels of RNA, we employed SMART-seq technology (Takara Biosciences) which can amplify cDNA directly from 1-1000 cells. Protocols were standardized using limiting serial dilution of cell line EV RNA (data not shown). EV RNA was isolated by EXOQUICK^®^ Exosome isolation and RNA purification kit for serum/plasma (System Biosciences,CA) and was reverse transcribed to make amplified cDNA by SMART seq v4 ultra-low input RNA kit (Takara Bio Inc, USA). After shearing of cDNA by Covaris ultrasound water bath sonicator and purification by Ampure beads (Beckman Coulter, Germany), a sequencing library was made using 1 ng of sheared cDNA using Low Input Library Prep Kit v2 (Takara Bio Inc, USA). DNA unique Dual index kit was used to combine libraries for sequencing (Takara Bio Inc, USA). NOVASEQ 3000 was used for sequencing at the NIBMG core facility (COTERI) to generate 25 million paired end reads per sample. Tru-cut biopsy samples were submitted to NIBMG core sequencing facility. Libraries were made using Illumina Truseq stranded sequencing kits and 50 million paired end reads were obtained per sample using NOVASEQ 3000.

### Data analysis

Upon receiving FASTA files, FASTQC check was performed to remove contrived data. FastQ files were trimmed by CutAdapt to remove the adapter sequences and reads were aligned with the human genome (GrCh38.1) by Hisat2.0 algorithm. Aligned sequences were then processed by Featurecounts package for acquiring transcript counts. TMM normalisation was performed to normalise the data and LIMMA VOOM package was used to obtain differentially expressed genes (DEGs) using Galaxy (www.usegalaxy.org). Z score calculation, spearman correlation analysis and heatmaps were generated in Bioconductor R using relevant packages. Pathway enrichment analysis was performed by EnrichR (https://maayanlab.cloud/Enrichr/). Venn diagrams were generated by Venny2.0. ROC analysis was performed by Graphpad Prism 8.4.

### Data availability

RNA sequencing data is available in GEO omnibus (GSE256523)

### TCGA data analysis

TCGA GDC portal was used to download TCGA Pan cancer data of Breast cancer, consisting of 1141 patients and 121 normal samples. STAR count files were downloaded and LIMMA was performed on the count file. List of DEGs at p<0.05, FC=1.5 were used to overlap with the EV-C set genes. DEGs from Firehose Legacy data with >100 control and > 700 tumor samples were obtained from Rosario *et al* (15).

### Quantitative real time polymerase chain reaction (qRT-PCR)

cDNA was amplified using SMART seq v4 ultra-low input RNA kit (Takara Biosciences). 0.5 ng of amplified cDNA from EV-H and EV-C RNA was used as template. SYBR green mix (KAPA BIOSYSTEMS, South Africa) was used for qRT-PCR, using primers described in Supplementary table 2. ΔC_T_ values (C_T_ Gene-C_T_ Actin) were calculated and plotted using Graphpad Prism 8.4.

### Statistical analysis

All the statistical analyses for transcriptomic data were conducted using Bioconductor R packages. Statistical analysis for qRT-PCR performed using GraphPad prism software. Comparisons between two groups were performed using student’s t-test and data was considered statistically significant at *p* ≤ 0.05. Significance in all figures is indicated as follows: * *p* ≤ 0.05, \*\**p* ≤ 0.01, \*\*\**p* ≤ 0.001 and \*\*\*\**p* ≤ 0.0001.

## Results

### Characterization of EVs

Quality control of EVs to analyse their size, shape, concentration was conducted by Nanoparticle tracking analysis (NTA) and immunoblotting for known protein markers in EVs (Figure 1A-B). NTA analysis showed enrichment of particles in a size range from100nm-300nm, with a mean size of 165nm (Figure- 1A). The concentration of EVs was calculated from the counts obtained after 3 repeat runs of the sample and the average count was 2X10^11^ EVs per ml of serum which is consistent with previously reported data. Immunoblotting of isolated EV proteins showed presence of known exosomal marker proteins CD81, CD63, TSG101, Flot1, Alix, GM130, ANAXA5, EpCAM and ICAM confirming that the isolated particles were enriched for exosomes (Figure 1B).

**Figure 1:**
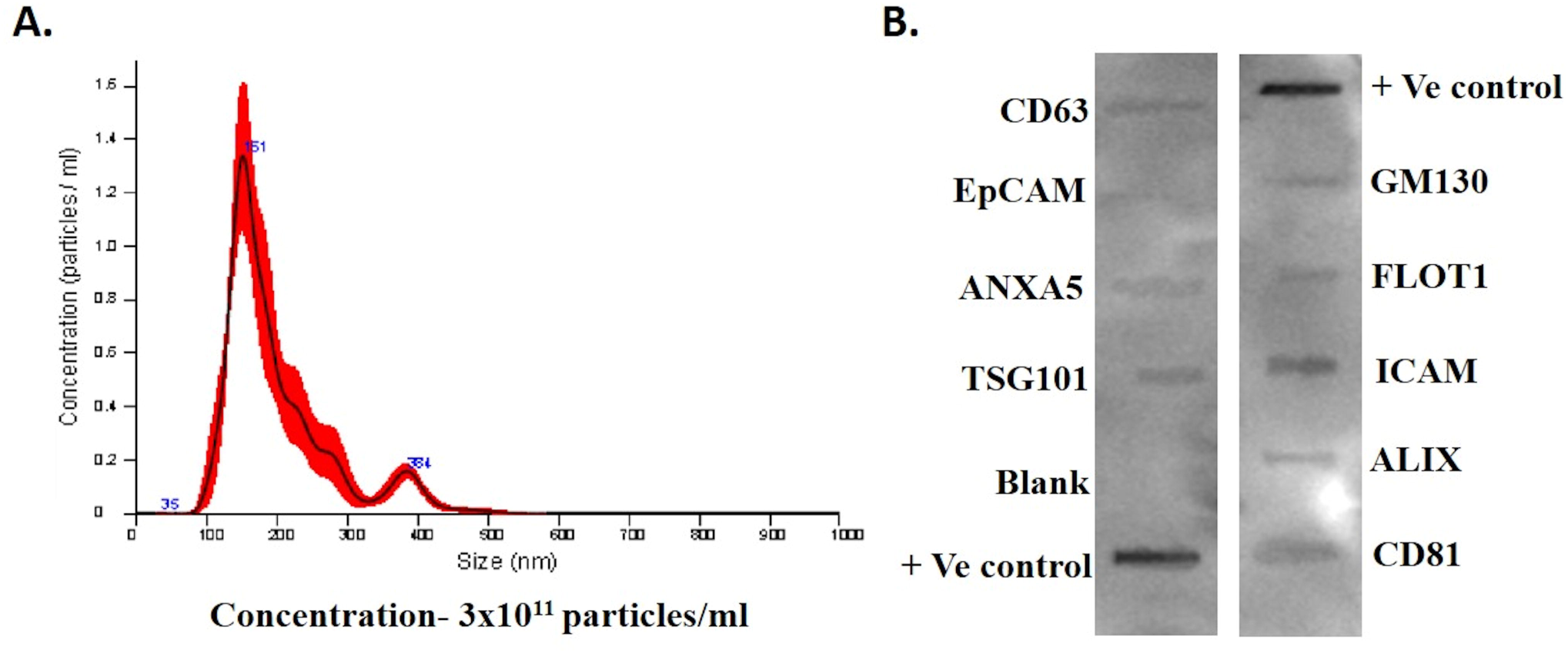
Characterization of EVs. A) Nano tracking analysis of serum derived EVs showing size and concentration. B) Immunoblot of known EV markers, positive and negative control

### EV RNAs may have the potential to diagnose cancer

We compared the transcriptome of serum derived EVs from breast cancer patients and healthy controls. Data was processed as described in methods. LIMMA-voom analysis identified 320 significant DEGs after multiple testing correction using Benjamini-Hochberg method (adjusted *p* ≤ 0.05) (Supplementary Figure 1A, Supplementary table 3). Interestingly, all DEGs found showed positive fold change expression in EV-Cs when compared to EV-Hs. These 320 DEGs (EV-C set) were able to clearly separate EV-C samples from EV-H as demonstrated by Z-score analysis of their normalized counts (Figure 2A). Pathway analysis showed enrichment in Myc target V1, PI3K/AKT/mTOR pathways, which have been shown to be enriched in multiple cell line EV datasets (11). In addition, changes in Heme metabolism, Interferon alpha and Interferon gamma, complement and Allograft rejection pathways were evident in our EV-C set (Figure 2B). Overall, the genes and their direction of expression indicated establishment of an immune suppressive environment.

**Figure 2:**
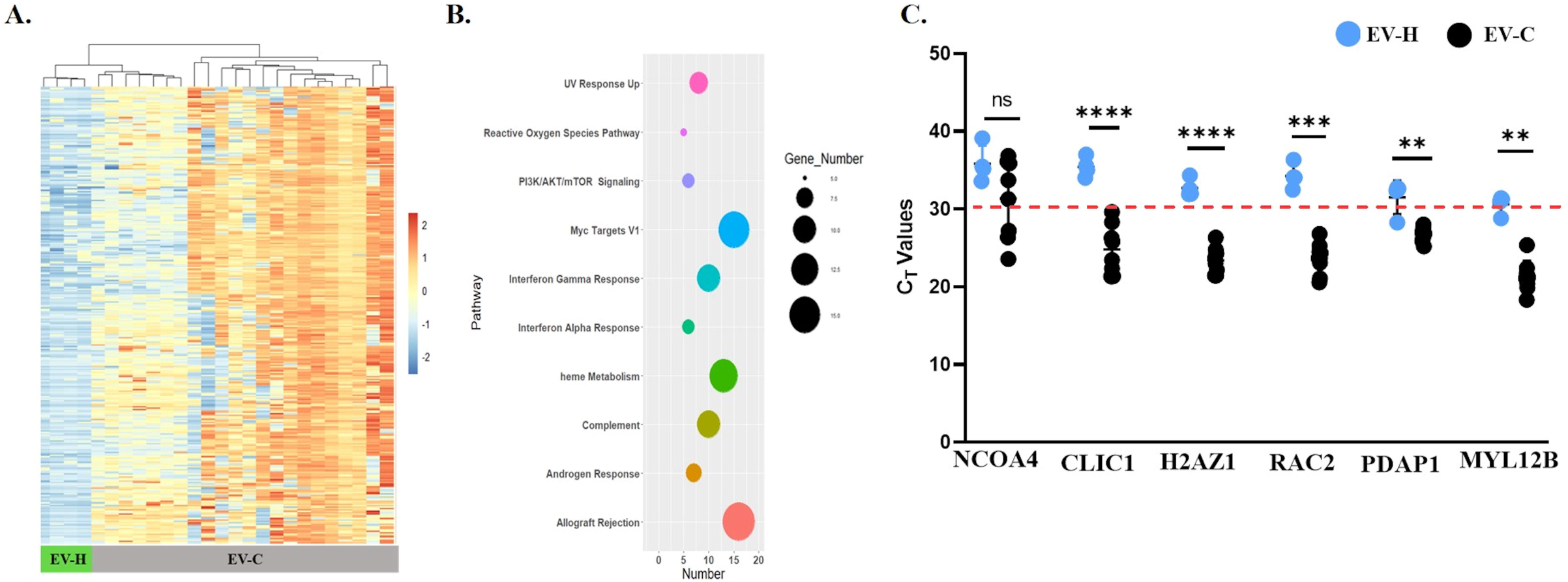
RNA Seq analysis of EV-C and EV-H samples. A) Z score-based clustering of EV set genes. B) Pathway enrichment analysis; C) qRT-PCR of 6 selected genes.

To determine if EV-C set genes could serve as discriminators between EV-C and EV-H samples, ROC analysis of top 15 significant DEGs was performed (Supplementary Figure 1B). All 15 genes had an area under the curve value (AUC) greater than 0.85, suggesting that they potentially, could classify cancer derived EVs from healthy sera EVs. Stripcharts obtained from LIMMA analysis showed consistently higher expression of all 15 genes in EV-Cs as compared to EV-Hs (Supplementary figure 1C). Based on literature and their prior involvement in breast cancer, we selected 6 genes with a stringent AUC cut-off of more than 0.9. CLIC1, RAC2, H2AZ1, MYL12B, PDAP1 and NCOA4 were further analysed and their expression levels in EV RNA-seq data are shown in Supplementary Figure 1D. qRT-PCR was performed with 4 EV-H and 8 EV-C samples to validate expression of these 6 genes (Figure 2C). The C_T_ values of 5 genes were significantly higher in EV-H samples as compared to EV-C, indicating that these genes had nearly undetectable expression in control samples but significant presence in EV-C samples. This observation validated the RNA-seq data. In contrast, NCOA4 expression was highly variable in both groups and therefore achieved no significance. Therefore, except NCOA4, the remainder subset of 5 genes expressed in EVs could be capable of effectively diagnosing breast cancer (Dia set).

### Probable origin of EV-C profiles

To understand origins of EV-C datasets, we evaluated two databases. First, we investigated if EV-C set could be classified on the basis of gene regulatory networks of unique transcription factors, since a primary regulator of gene expression in breast cancer is ER. CHEA3 analysis that represents individual ChIP-seq, ENCODE and literature-based surveys of TF binding sites, was employed to explore the EV-C set. In circulating EVs, we did not find ER regulated genes, but identified SPI1 as the major TF for 159 out of 320 genes. 4/5 Dia set genes described in the previous section are SPI1 targets. SPI 1 RNA itself was also recognized as one of the DEGs (Figure 3A). ROC analysis showed it did not possess an AUC value as high as Dia set and PCRs were not conducted for this gene (Supplementary figure 2). The second most abundant TF was NFE2, and both NFE2 and SPI1 regulate each other. Incidentally, expression of SPI1 is enhanced in osteoclast and areas of the bone prior to metastatic spread of tumor cells (16) and our data implies that SPI1-target genes in EVs could influence this niche.

**Figure 3:**
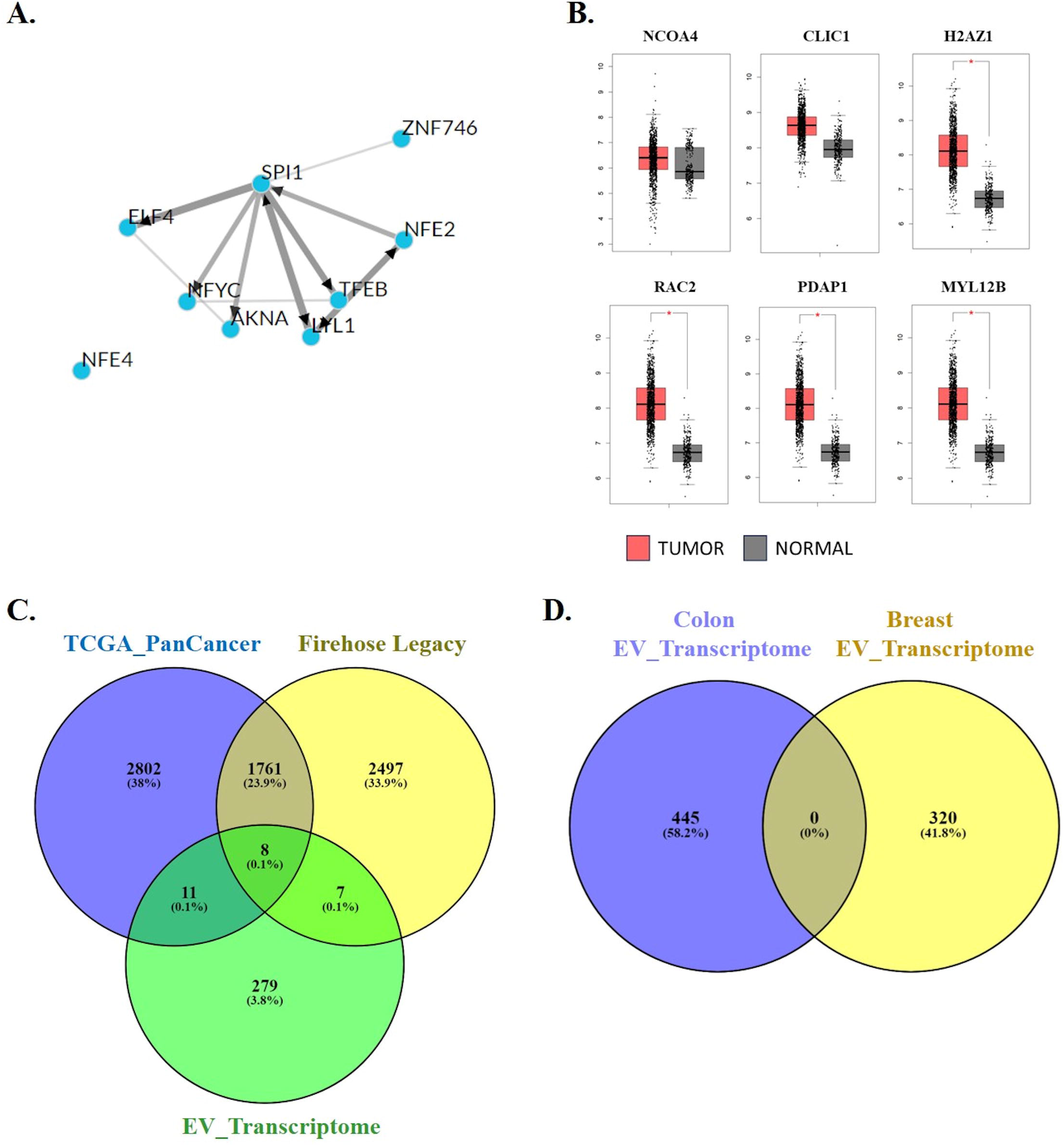
Analysis of EV set genes with other cancer datasets. A) ChEA3 data showing interactome of potential regulatory transcription factors for EV set genes. B) Expression of 6 selected genes in breast cancer and normal samples. C) Overlap between EV

The second approach we used was to explore expression of select genes in breast cancer datasets. We interrogated the GEPIA data and four genes, except NCOA4 and CLIC1, were highly expressed in tumor as opposed to normal samples (Figure 3B). As seen in our EV transcriptome and qRT-PCR data, NCOA4 expression was variable both tumor and normal samples. This encourages us to estimate the extent of overlap between the changes in gene expression in circulating EVs versus those found in breast cancer tumors. Analysis of two public TCGA datasets was conducted as described in methods. 4563 DEGs (adjusted *p* value<0.05, FC-1.5) were obtained from TCGA Pan cancer data by comparing breast cancer tissue RNA profiles with normal healthy tissue. Only 19 genes overlapped with the 320 EV-C set genes. Upon similar analysis of Firehose Legacy data, 4258 DEGs were obtained and 15 genes were common with the EV-C set (Figure 3C). Overall, we were unable to find high tumor profile representation in EV transcriptomic data. Upon ROC analysis, this overlapping set of genes did not show AUCs more than 0.65 and were not explored by qRT-PCR. It is possible that EVs illustrate a systemic response to tumor presence rather than overall tumor gene expression profiles. Further, the other circulating EV dataset is published in colon cancer and analysis similar to ours has been carried out (17). The EV-C set obtained appears to be breast cancer specific because when compared to the study on colon cancer, not a single gene was common between the two sets (Figure 3D).

### EV transcriptome and ER status

Dichotomy of gene expression profiles based on ER status of breast tumors has been firmly established. We studied if EV-C RNA profiles reflected these differences. Comparison of EV-C transcriptome was carried out by assigning ER+ and ER- patients to separate groups (Supplementary table 4). First, Spearman’s Rank correlation analysis was done using all genes (21,000 features in RNA-seq data) expressed in the EV-C samples. Correlation map showed that both EV-C-ER+ and EV-C-ER- samples had better intra-group rather than inter-group correlation, with EV-C-ER+ intragroup coefficients being higher than those of ER- group (Figure 4A). This observation promoted the notion that serum EV profiling might have the potential to separate tumors based on ER status. 374 DEGs (p <0.001, logFC=2) were identified between EV-C-ER+ and EV-C-ER- samples. These genes are designated as the EV-C-ER set (Supplementary Figure 3A). Here stringent multiple hypothesis testing recovered a handful of genes due to low sample numbers in each group. Nevertheless, Z score was calculated from normalised counts data and the heatmap of EV-C-ER clusters is shown in figure 4B (Figure 4B). EnrichR identified Myc target V1, Adipogenesis, DNA repair, Inflammatory response, IL-6/JAK/STAT signalling and others as significantly enriched pathways in these subsets of genes (Figure 4C). Interestingly, classification of ER+ versus ER- breast cancer using serum EVs was not limited to ER responsive genes, since E2 response pathway genes were not detected in circulating EVs. These data indicate that additional genes profiled in EVs could be useful in segregating ER+ versus ER- tumors. ROC analysis of the top 15 genes obtained in this list is shown in Supplementary figure 3B. Most genes had AUC >0.75, indicating that these need to be validated in larger cohorts before they can be used to segregate ER- from ER+ tumors based on EV characterization.

**Figure 4:**
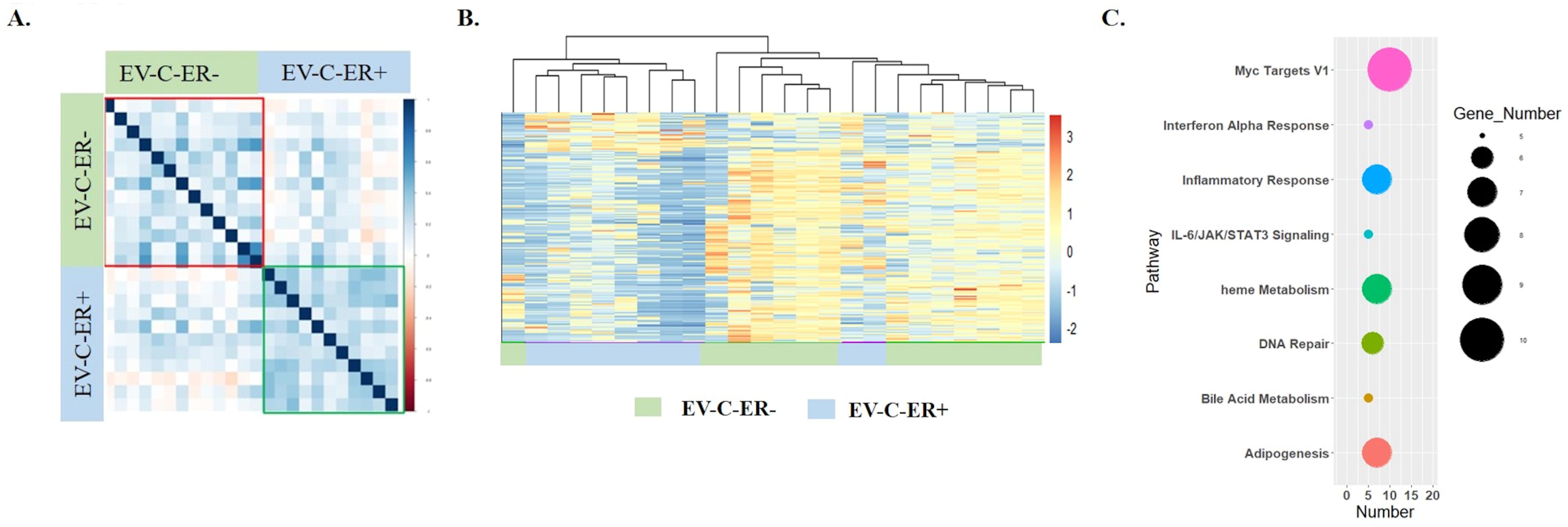
Analysis of EV-C-ER+ and EV-C-ER- samples. A) Spearman correlation of EV- C-ER+ and EV-C-ER- samples. B) Z score of normalised counts of 374 DEGs. C) Pathway enrichment analysis of 374 DEGs

For the EV serum samples studied, we had obtained matched tru-cut biopsy samples and profiled them by RNA-sequencing. Since, adjacent normal samples were unavailable, we segregated tumor profiles based on their ER status and identified a total of 850 DEGs (adjusted *p* value <0.05). Comparison with earlier reports showed significant reproducibility with previously described ER target genes (data not shown). When compared with the EV-C- ER list, only 11 genes overlapped with the tumor DEGs (Supplementary figure 3D). This result supports our idea that circulating EVs show a distinct set of genes that could be useful in binning diagnosed cancer into ER+ versus ER-.

### EV transcriptome can separate node negative and metastatic tumors

Our tumor samples included 7 patients with early disease without metastasis to lymph nodes or distant sites, (EV-C-NM) and 9 women with advanced disease and distant metastases (EV- C-M). Spearman’s Rank correlation analysis demonstrated that while the EV-C-NM group had high correlation in gene expression within the group, the EV-C-M were quite distinct within themselves (Figure 5A). 640 DEGs that cluster these 16 samples into 2 groups are shown in Figure 5B (Supplementary figure 4A, Supplementary table 5). Pathway enrichment analysis indicated the PI3/AKT/mTOR signalling as most significant, a pathway already known to have a role in breast cancer metastasis (Figure 5C). Other significant pathways included Apical junction, IL6-JAK-STAT and Oxidative phosphorylation. Our earlier CHEA3 analysis of EV-C set had identified SPI1 and its targets that could be involved with metastatic niche formation. Interestingly, the same TF appeared in the EV-C-NM vs EV-C-M datasets strengthening the hypothesis that EVs may carry its targets and they may impinge on preparation of metastatic regions. However, the target genes identified here were distinct from those in EV-C set indicative of a vast and prominent SPI1 target RNA cargo in circulating EVs from breast cancer patients. In addition, the general transcription factor, GTF2B was also indicated.

**Figure 5:**
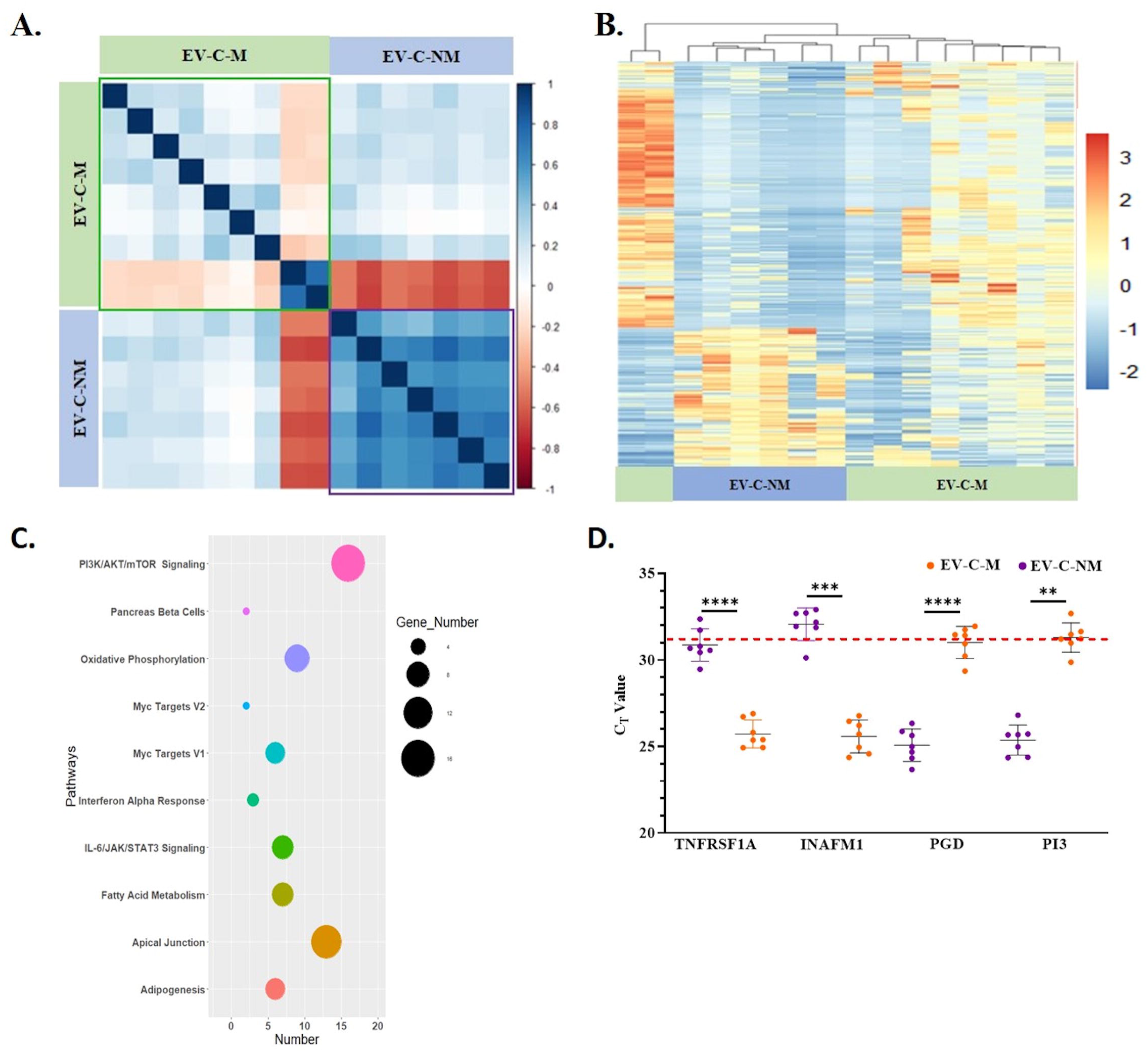
Analysis of EV-C-NM and EV-C-M samples. A) Spearman correlation of EV-C- NM and EV-C-M samples using normalised counts; B) Z score clustering of 640 differentially expressed genes. C) Pathway enrichment analysis. D) qRT-PCR of 4 selected genes from EV-C-M and EV-C-NM

ROC analysis of the top 15 genes was performed (Supplementary figure 4B). Four genes, PI3, PGD, INFM1, TNFRSF1A, having high area under the curve (AUC > 0.9), and prior relevance to cancer, were selected as the Met set. Strip charts of these genes are shown in Supplementary figure 4C). PI3 and PGD were highly expressed in EV-C-M samples whereas INAFM1 and TNFRSF1A were higher in EV-C-NM. As shown in figure 5B, these 4 genes faithfully clustered metastatic versus non-metastatic samples. The selected 4 genes were further validated by qRTPCR among 4 EV-C-NM and 4 EV-C-M samples (Figure 5D)

### Validation of select genes in an independent sample cohort

We collected additional 14 samples blinded for their cancer status and performed qRTPCR analyses for the 6 EV-C set and 4 Met set genes (Figure 6A). We could correctly classify healthy versus cancer samples each time confirming that these EV genes had a robust diagnostic potential. These genes could form the basis of identifying an EV-based cancer signature. Additionally, we engaged 11 more samples in qRTPCR analyses that were blinded for their metastasis status (Figure 6B). Here, we could correctly classify samples only 60% of the time when all 4 genes were employed. Interestingly, TNFRSF1A gene alone could consistently distinguish between metastatic and non-metastatic samples. Since sample numbers in our study are limiting, a separate study with a larger cohort of samples is essential to conclude the veracity of these genes as metastatic indicators.

**Figure 6:**
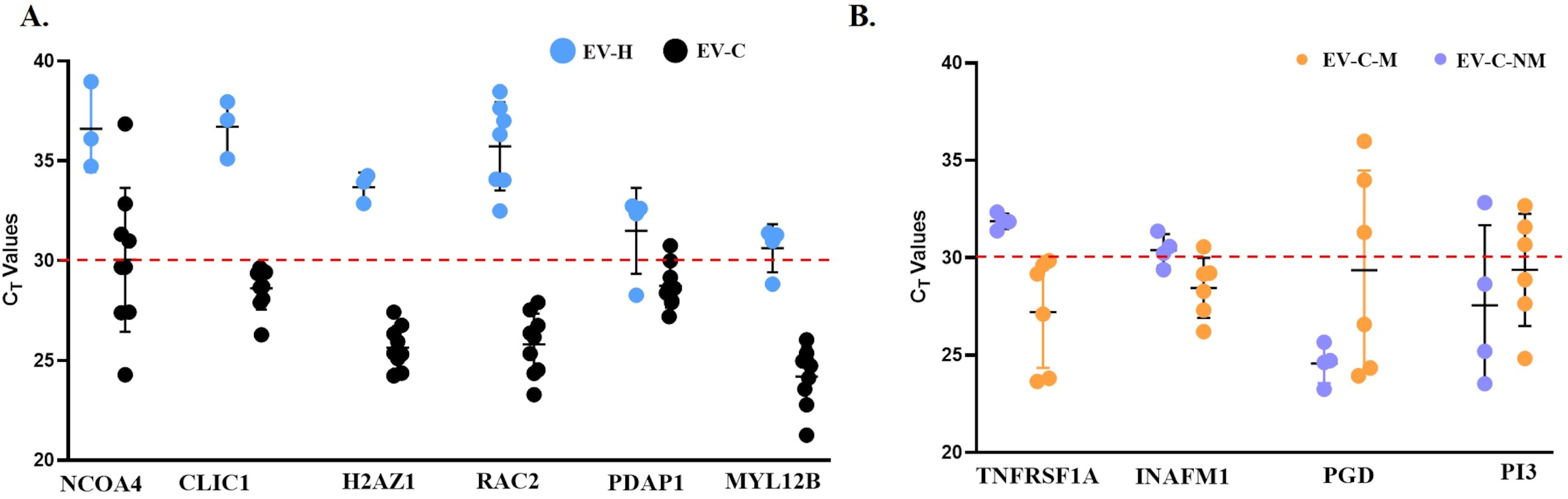
PCR on validation cohort. qRT-PCR for 14 blinded samples for A) 6 selected genes. B) 4 selected genes to distinguish metastatic and non-metastatic samples

## Discussion

Apart from three recent attempts using Ncounter platform (14) and a colon cancer study (17) and a study on long RNA of circulating EV from breast cancer (18) there are very few published studies on serum EV total RNA-seq analysis, due to lack of suitable and reproducible protocols. We established a protocol using 500µl of serum for EV RNA isolation followed by using Smart-seq V4 ultra-low input RNA kit for cDNA amplification and subsequent transcriptome library synthesis. First, we compared transcriptome profile of EV-C and EV-H individuals to obtain 320 DEGs that successfully clustered normal and tumor groups separately. This implies EVs in circulation may carry a vastly different RNA cargo between cancer and disease-free samples. More than 50% of the RNAs found appear to be targets of SPI1/NFE2 transcription factors. These factors are known to be expressed in areas of bone that homing of tumor cells (16,19). Pathway enrichment analysis showed predominance of immune suppressive indicators such as reduced Allograft rejection, Interferon gamma (IFNγ) and Interferon lambda (IFNλ) activity, which supported the overall propensity of tumor-induced immune evasion, a hallmark pathway in cancer (2). Both qRT-PCR based testing of RNA-seq samples and EV samples from validation cohorts blinded for presence or absence of breast cancer indicated that 5 genes, CLIC1, RAC2, H2AZ1, MYBL12B and PDAP1, accurately classified cancer and healthy samples. All have been implicated in tumor progression earlier and showed higher expression in tumors when compared to normal samples in public datasets (20–22). However, in our analysis of 320 genes with TCGA BRCA RNA-seq profiles, only 19 genes overlapped. In colon cancer, 58/450 genes were found to overlap with colon cancer tumor dataset DEGs (17). No overlap was found between the circulating EV profiles of colon and breast cancer. It is possible that circulating EVs are not phenocopies of tumors they originate from and may display organ specific, yet profiles that illustrate systemic change due to presence of a tumor. Similar conclusions have been drawn earlier in colorectal cancer and mass spectrometric analysis of EVs in breast cancer (23,24).

Comparison of EV-C-ER+ and EV-C-ER- samples showed 372 genes, most were not ER targets but genes in the IL6-JAK-STAT, inflammatory response, interferon pathways previously known to be differentially altered in ER+ versus ER- tumors (25). IL6 JAK-STAT pathway plays a significant role in TNBC metastasis raising the possibility that these RNAs are carried via EVs. Earlier data shows that EVs establish organotropism by using surface integrins as ‘zipcodes’ and also the premetastatic niche (26). We were curious if comparison of EV-C-NMs with EV-C-Ms would yield any DEGs that could discriminate between the two groups. Amongst others, pathway enrichment analysis of 640 genes showed PI3-AKT-mTOR pathway as most significant pathway, which is known to influence metastasis of ER+ breast cancers (27). Genes bearing 4 highest AUC values were able to cluster of M and NM tumors according to their gene expression pattern in RNA-seq samples and these levels could be validated by qRT-PCR analysis of serum EV RNAs. However, in the validation set of 14 samples, only TNFRSF1A was able to accurately classify NM versus M samples. An 8 protein EV signatures has been proposed earlier establishing the capacity of EVs to monitor and assess response to chemotherapy in MBC patients(28). However, single marker did not have high sensitivity and specificity. In our data, a single RNA appears to be capable of classification. However, exploring a larger cohort of patients for TNFRSF1A RNA expression in EVs for detecting metastatic status is necessary to prove this hypothesis.

Despite the limitation of a small sample size, our study provides an analysis of mRNA profiles of circulating EVs and identifies cancer relevant genes and pathways in Breast Cancer patients (EV-C) that are absent in EVs from healthy female sera (EV-H). The DEGs were assayed by qRT-PCR analysis, and EV RNAs that could potentially diagnose breast cancer, distinguish ER positive and /ER negative samples and segregate metastatic and non-metastatic samples were identified. A minimum set of 5 Dia set and 1 Met set gene(s) have been validated using samples blinded for their tumor status. Our data lays the foundation for identifying such EV-based RNA markers of disease states in breast cancer. Whether EVs are better than other components of liquid biopsy, namely circulating cell free DNA (ctDNA) or circulating tumor cells (CTCs), remains to be studied in matched samples in a clinical trial setting.

## Conclusion

Liquid biopsy is a promising approach to diagnose and monitor disease progression and response to therapy. We have profiled circulating extracellular vesicles obtained from healthy controls and breast cancer patients. Quantitative RT-PCR of a 5 -gene EV panel derived from transcriptomic data was used successfully to accurately discriminate healthy and cancer samples from a validation cohort. EV RNA cargo substantially differed between ER+ and ER- tumors and between metastatic and non-metastatic states. Our data pin-points genes and pathways that could aid detection and classification of breast cancer using circulating EVs.

## Declarations

### Ethics Approval and Consent to participate

This study was approved by the Tata Medical Center, Kolkata Institutional review board (IRB EC/PHARMA/34/17).

### Availability of data and materials

Data is available in GSE 256523

### Authors contributions

**A Gupta**: Data generation, visualization, analysis and writing of first draft, review and editing of final draft of manuscript. **R. Ahmed**: Collection, clinical data generation, review and editing **S. Agarwal**: Collection, clinical data generation, review, and editing. **G. Mukherjee**: Clinical Histopathology, review and editing **KV Desai**: Conceptualization, Supervision, Project coordination, data generation, analysis and writing of first draft, review and editing of final draft of manuscript

## Acknowledgement

All the patients who have participated in the studies are duly acknowledged along with Hospital staff. NIBMG sequencing facility is acknowledged for extending use of their instrumental facility.

## Abbreviations

EV: Extracellular vesicles
CTCs: Circulatory tumor cells
cfDNA: cell free DNA
LncRNA: Long non-coding RNA
mRNA: messenger RNA
DEGs: Differentially expressed genes
AUC: area under the curve
ER: Estrogen Receptor
AR: Androgen receptor
NTA: Nanoparticle tracking analysis
EV-C: Extracellular vesicles from cancer patient
EV-H: Extracellular vesicles from healthy individuals
EV-C-ER+: Extracellular vesicles from ER+ cancer patient
EV-C-ER-: Extracellular vesicles from ER- cancer patient
EV-C-M: Extracellular vesicles from metastatic patient
EV-C-NM: extracellular vesicle from non-metastatic patient
qRT-PCR: quantitative Real time Polymerase Chain Reaction
IFN: Interferon
OxPhos: oxidative phosphorylation

## Figure Legends

**Supplementary Figure 1:** A) Volcano plot of EV set genes obtained by LIMMA of EV-C and EV-H RNA-seq data. B) ROC analysis of top 15 DEGs. C) Strip charts of top10 DEGs. D) Z score-based clustering of 6 selected genes

**Supplementary Figure 2:** ROC analysis of SPI1

**Supplementary Figure 3:** A) Volcano plot of DEGs obtained by LIMMA of EV-C-ER+ and EV-C-ER- RNA-seq data. B) ROC analysis of top 15 DEGs

**Supplementary Figure 4:** A) Volcano plot of DEGs obtained from LIMMA of EV-C-M and EV-C-NM RNA-seq data. B) ROC analysis of top 15 DEGs. C) Strip charts of top10 DEGs. D) Z score-based clustering of 4 selected genes

**Supplementary table 1:** Patient information

**Supplementary table 2:** Oligonucleotide sequences of RT-PCR primers

**Supplementary table 3:** EV set genes

**Supplementary table 4:** DEGs from EV-C-ER- and EV-C-ER+

**Supplementary table 5:** DEGs from EV-C-M and EV-C-NM

