## Supplementary figures and images for "Exploiting gene expression profiles of circulating extracellular vesicles for breast cancer detection"

### Supplementary Figure 1

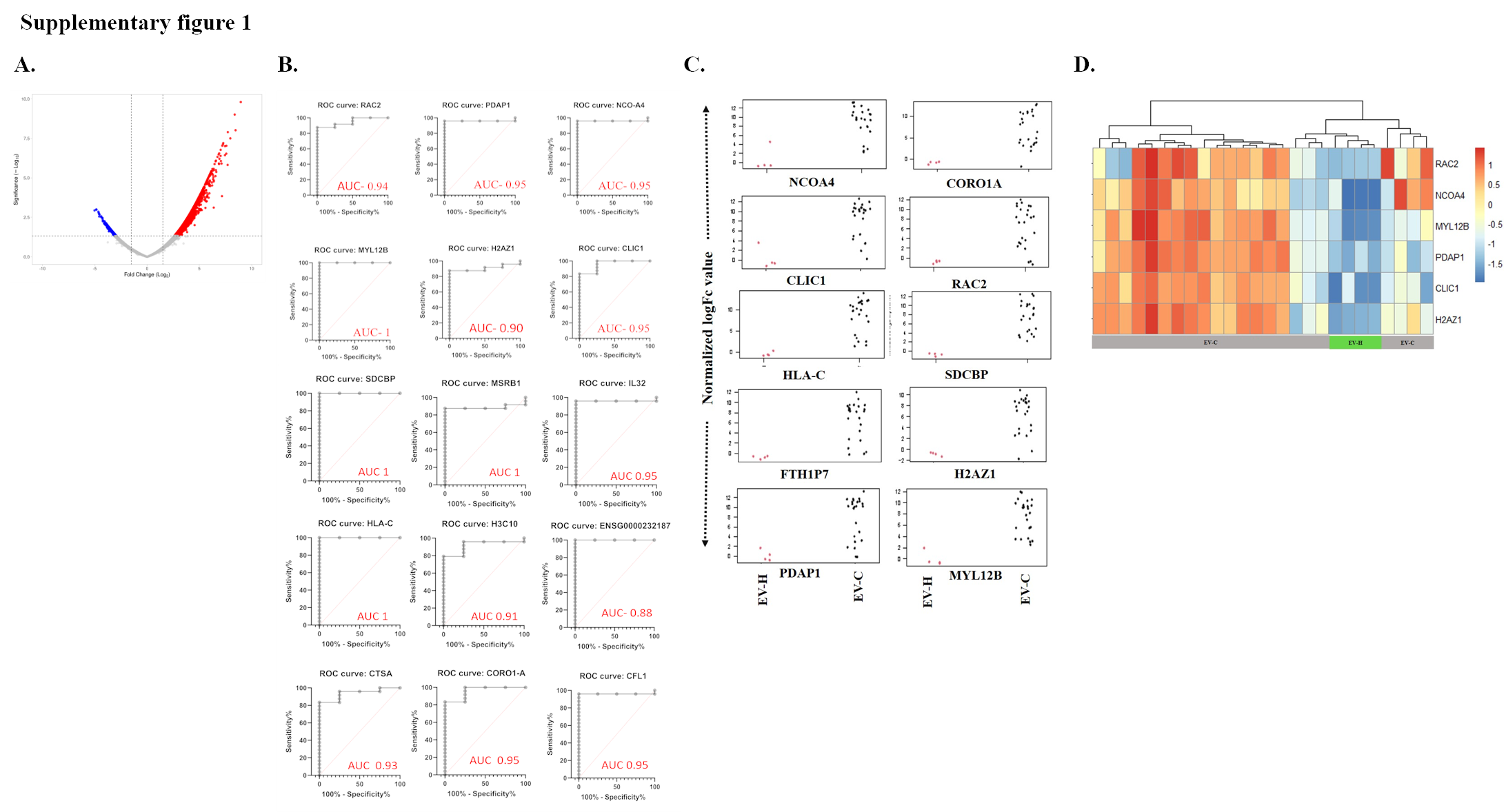

### Supplementary Figure 2

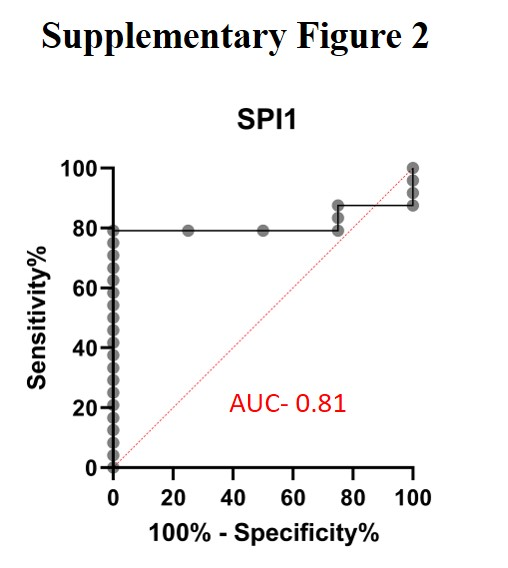

### Supplementary Figure 3

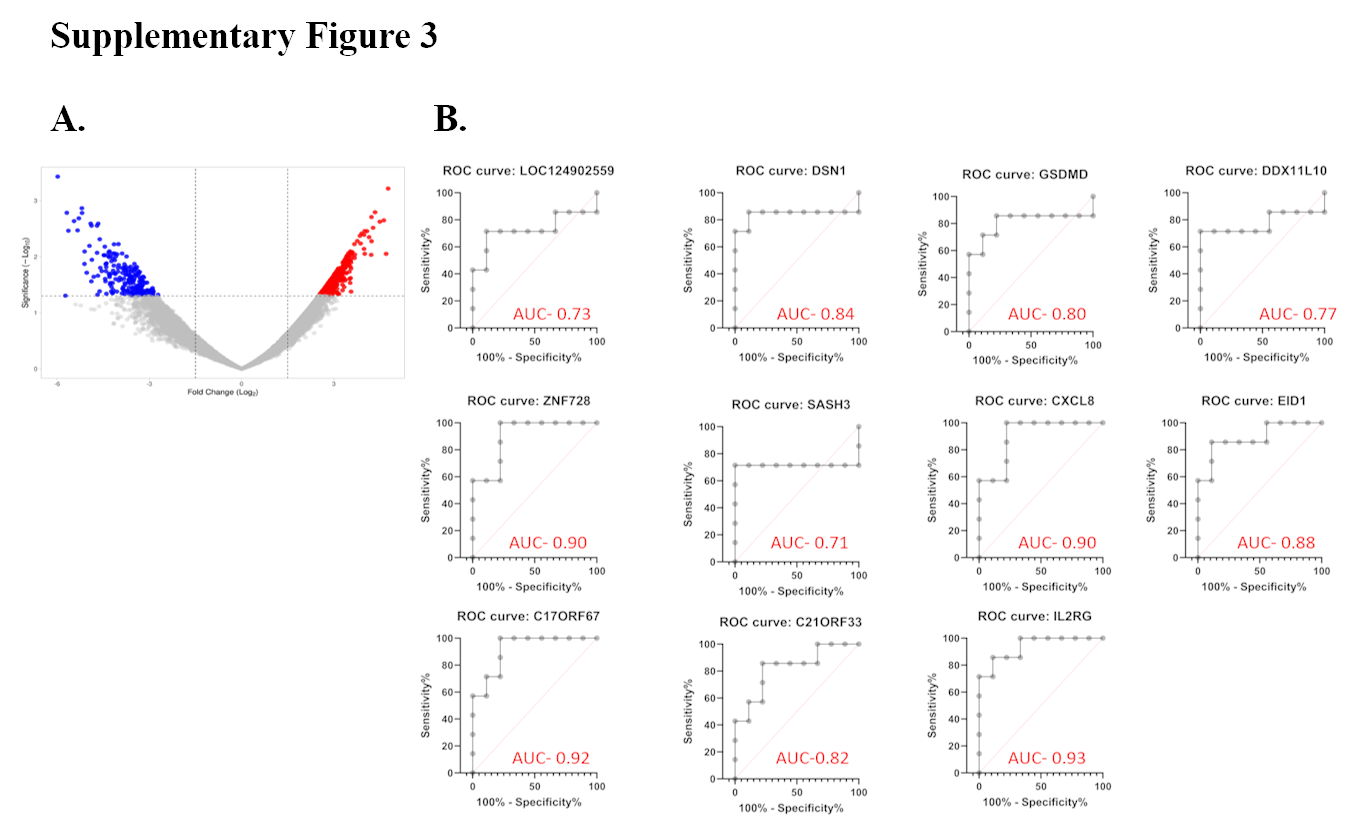

### Supplementary Figure 4

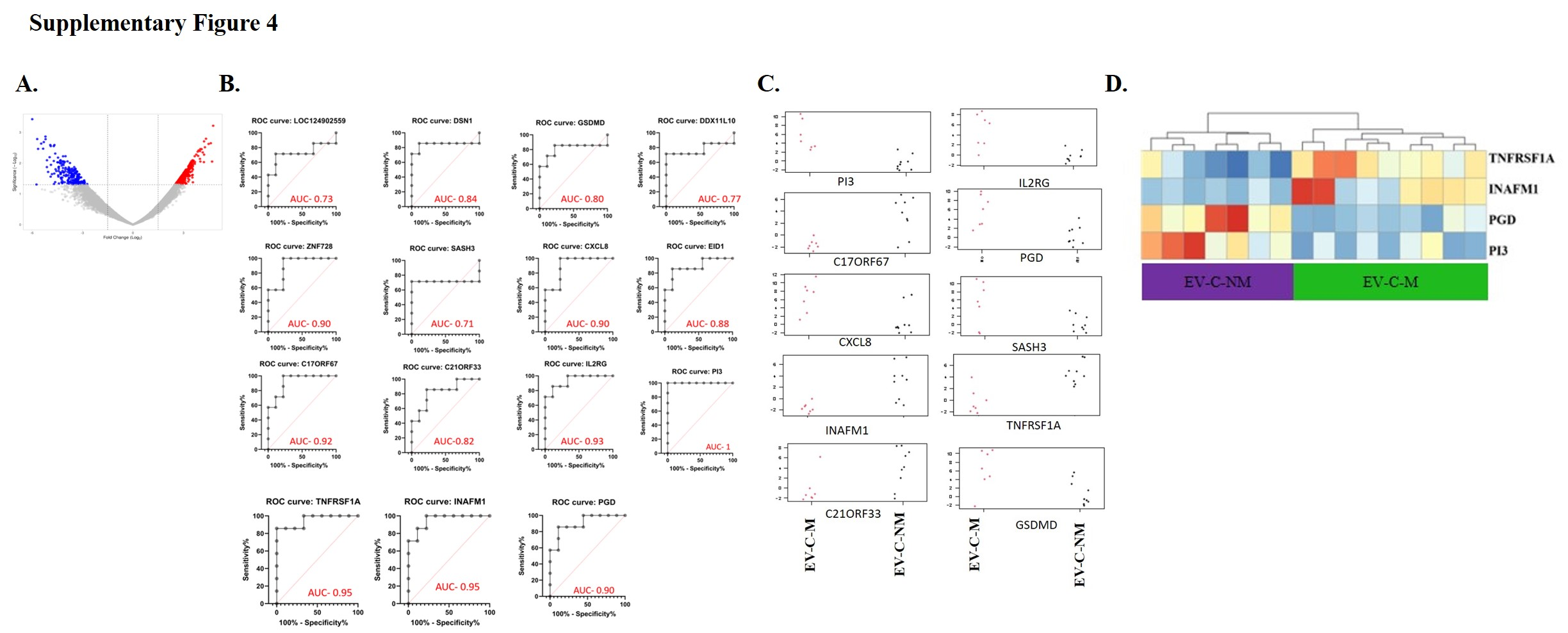
